# Learning with filopodia and spines

**DOI:** 10.1101/2023.08.26.554942

**Authors:** Albert Albesa-González, Claudia Clopath

## Abstract

Filopodia are thin synaptic protrusions that have been long known to play an important role in early development. It has recently been found that they are more abundant in the adult cortex than previously thought, and more plastic than spines (button-shaped mature synapses). Inspired by these findings, we introduce a new model of synaptic plasticity that jointly describes learning of filopodia and spines. The model assumes that filopodia exhibit additive learning, which is highly competitive and volatile. At the same time, it proposes that if filopodia undergo sufficient potentiation they consolidate into spines, and start following multiplicative learning dynamics. This makes spines more stable and sensitive to the fine structure of input correlations. We show that our learning rule has a selectivity comparable to additive spike-timing-dependent plasticity (STDP) and represents input correlations as well as multiplicative STDP. We also show how it can protect previously formed memories and act as a synaptic consolidation mechanism. Overall, our results provide a mechanistic explanation of how filopodia and spines could cooperate to overcome the difficulties that these separate forms of learning (additive and multiplicative) each have.

**Author Summary:** Changes in the strength of synaptic connections between neurons are the basis of learning in biological and artificial networks. In animals, these changes can only depend on locally available signals, and are usually modeled with *learning rules*. Based on recent discoveries on *filopodia*, a special type of synaptic structure, we propose a new learning rule called Filopodium-Spine spike-timing-dependent-plasticity. Our rule proposes that filopodia follow additive STDP and spines (mature synapses) multiplicative STDP. We show that our model overcomes classic difficulties that these learning rules have separately, like the absence of stability or specificity, and can also be seen as a first stage of synaptic consolidation.

## Introduction

*Filopodia* are thin protrusions in dendrites (Jontes & Smith, 2000) that have been long-known to exist. Until recently, though, they were thought to play an important role only at developmental stages of brain formation (Fiala et al., 1998). However, filopodia have been now found to be more abundant than previously thought, as well as the structural substrate for silent synapses in the adult brain (Vardalaki et al., 2022). In fact, these structures conform to around 30% of the dendritic protrusions. Because they lack AMPA channels, these synapses are effectively silent, meaning they cannot elicit a postsynaptic response unless the postsynaptic neuron is depolaraized. Furthermore, filopodia are sensitive to plasticity induction protocols that are insufficient for spines (i.e., mature synapses that do contain AMPA channels). In particular, when applied the same spike-timing-dependent potentiation protocol, filopodia increased their synaptic efficacy but spines’ remained the same. Finally, within minutes of being potentiated, filopodia’s appearance started resembling that of a spine (Vardalaki et al., 2022). These findings beg the question: what are the underlying learning mechanisms of filopodia, and how do they coordinate with spines to facilitate cognitive functions? While there have been previous proposals of the distinct functional roles of filopodia and spines (Ozcan, 2017), experimental evidence that supports these has been scarce, and their relation to computational models of synaptic plasticity unexplored.

The protocol used in Vardalaki et al. (2022) is inspired by early studies of *spike-timing-dependent plasticity* (STDP) (Bi & Poo, 1998). In STDP, changes in the weights’ strength are a function of the difference in timing between pre and postsynaptic neurons. Two prominent computational models of STDP are additive (add-STDP) (Gerstner et al., 1996; Song et al., 2000) and multiplicative (mult-STDP) (Van Rossum et al., 2000). Add-STDP yields highly selective, bimodal receptive fields, making it a compelling learning rule to uncover salient statistical patterns in the input structure. However, it is intrinsically unstable, requires imposing hard-bounds on the weights, and is only able to capture the coarse structure of input correlations. On the other hand, mult-STDP creates a weight distribution that continuously matches the correlation structure of presynaptic input. These unimodal distributions are typically considered more realistic, since they better reproduce experimental results. However, the absence of synaptic specialization can hinder learning by mapping very different patterns to the same neuronal output activity. How these two different pictures can be reconciled has puzzled neuroscience modelling research for years (Morrison et al., 2008). One hypothesis is that the unimodal distributions found in experiments are in fact only the *observable* part of all synapses. These would be complemented with a big proportion of *silent synapses* (Brunel et al., 2004; Barbour et al., 2007), which are not detectable via changes in postsynaptic potential, and would form another pool of effectively silent synapses. Whether these putative silent synapses were in fact present remained, until now, largely unclear.

In this work, we present a computational model that explicitly distinguishes between filopodia and spines (Filopodium-Spine STDP, FS-STDP). We hypothesize that filopodia learning can be well described by add-STDP, while spines learn in a multiplicative manner. As suggested by experiments (Vardalaki et al., 2022), filopodia that undergo potentiation can be converted into spines, and our model assumes that the inverse is also possible. We turn to previous results of additive and multiplicative learning to predict the functional advantages of this type of combined learning, and use simulations to confirm our hypothesis. In particular, we show that FS-learning establishes a two-stage competition. The first stage is strongly competitive and classifies synapses in a bimodal fashion (as does add-STDP) depending on presynaptic correlations. The second stage, governed by mult-STDP, continuously represents the input correlations in the spines that emerge from the first stage. We also show that the soft-bounded dynamics of mature spines shield them from becoming filopodia again. This makes the gross structure of the existing receptive field resistant to new correlated input, thus protecting previously formed memories from being erased instantaneously when environmental statistics change.

## Results

### FS-learning induces strong competition between filopodia and weak competition between spines

Our model considers the existence of *filpodia* (Figure 1A, in grey), which can dynamically evolve through learning into *spines* (in purple). We hypothesize that filopodia, which are more sensitive to potentiation protocols, follow additive dynamics. In additive STDP, the changes in weights depend exponentially on the differences of spike times between the pre and postsynaptic neuron. Furthermore, it is independent of the synaptic state, so it is not affected by the current strength of the weight (Figure 1B, Top). As has also been found in experiments (Vardalaki et al., 2022), we assume that if a filopodium is potentiated for a long enough time, it is converted into a spine (Figure 1A). We further propose that spines, in contrast, follow multiplicative learning. Mult-STDP incorporates soft-bounds that make weight changes dependent on the synaptic state (Figure 1B, Bottom). This form of learning is intrinsically more stable, has been experimentally reported in spines (Montgomery et al., 2001; Loebel et al., 2013), and is also consistent with Vardalaki et al. (2022), where it was harder to modify the synaptic efficacy of spines compared to filopodia.

**Figure 1.**
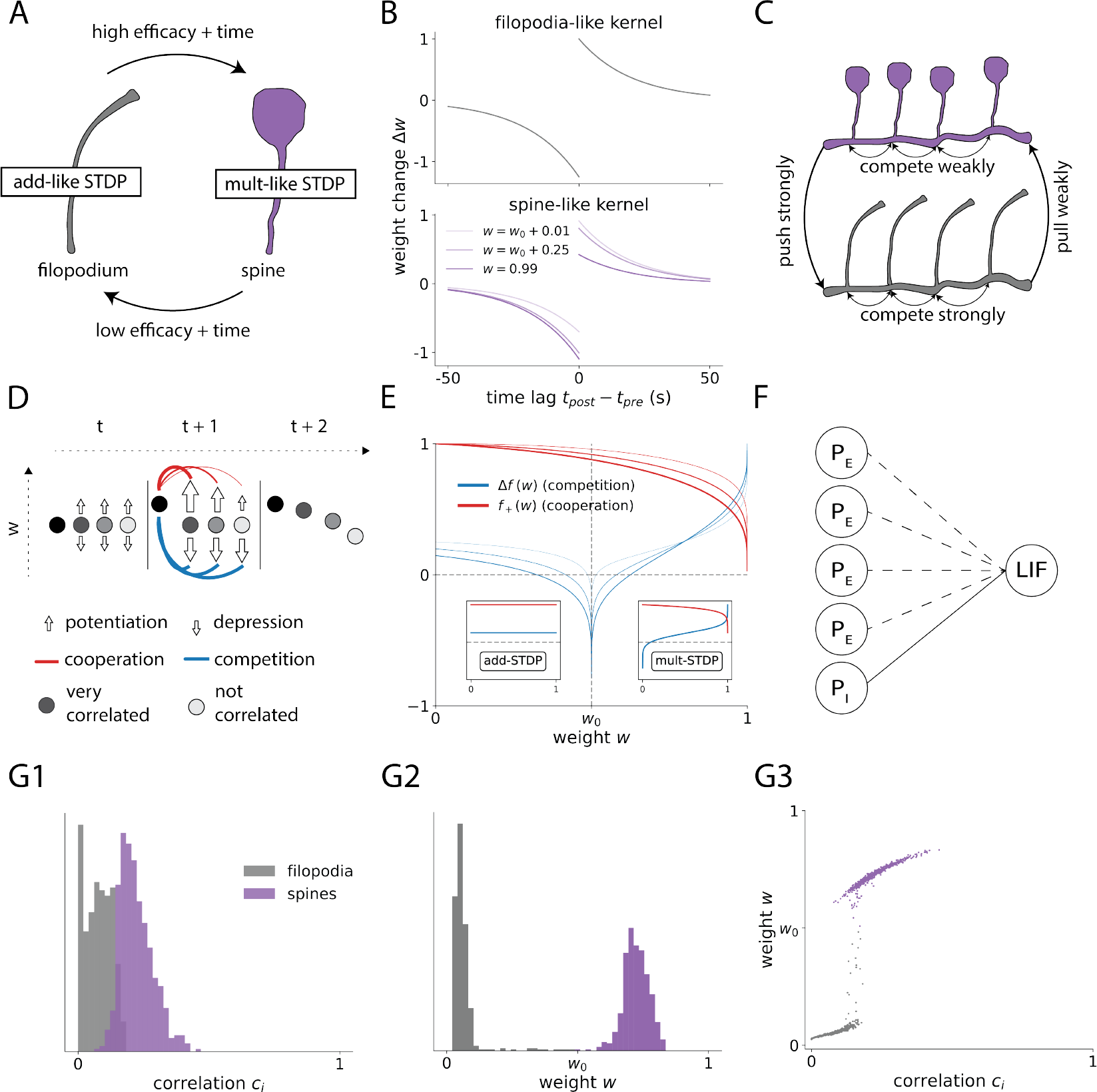
Filopodia compete strongly to become spines, and spines compete weakly to represent input correlation. **A:** Filopodium (grey) -Spine (purple) scheme. Filopodia can become spines if their efficacy is increased for a long enough time (observed experimentally in Vardalaki et al. (2022)). Similarly, spines can become filopodia if their weight is consistently low (our model). **B:** Learning for filopodia (Top) and spines (Bottom). Filopodium-like dynamics effectively lead to additive learning, and spine-like dynamics to multiplicative learning. **C:** Diagram indicating the push-pull forces between the different types of synapses in our model. Spines are strong weights and compete weakly between them. Filopodia are silent synapses that compete strongly to become spines. While the push that spines exert over filopodia is strong, the pull that filopodia exert over spines is weak (due to the lower-soft bound *w*_0_ in the dynamics of spines). **D:** Diagram of competition (blue lines) and cooperation (red lines). When a synapse increases its efficacy (left circle from t to *t* + 1), it has two opposite effects on the rest of the synapses. It equally increases depression (competition) and it also increases the potentiation of those it is correlated with (cooperation). **E:** Competition (blue) and cooperation (red) terms, as a function of weight w, under equilibrium conditions. *w*_0_ is a model parameter that sets the lower soft-bound of spines. For this reason, spines are distributed between *w*_0_ and 1. This is again used to classify a weight as filopodia (*w*_0_ < 0) or spine (*w*_0_ ≥0). Small subpanels on the left and right show, respectively, how the same plot would look for add-STDP and mult-STDP. The shape of competition and cooperation is very similar to additive learning for weights below *w*_0_, but resembles multiplicative learning for efficacies above this value. Thicker lines indicate higher values of the protection parameter a (Methods, Equation (4)). Higher values of a make transitioning back from spine to filopodia harder (competition becomes negative -cooperative-around *w*_0_). **F:** Diagram of network architecture. Our model consists of one postsynaptic LIF neuron that receives many Poisson realizations of presynaptic input, which can be either excitatory (*P*_*E*_) or inhibitory (*P*_*I*_). Only excitatory synapses are plastic. Excitatory presynaptic neurons are temporally correlated via a *correlation structure* 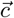 The spike times of presynaptic neurons *i* and *j* will be correlated only if both *c*_*i*_ and c_*j*_ are sufficiently high 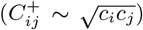. **G:** Example of FS-learning imprinting a pattern with correlation structure 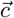 **G1:** Distribution of correlation strength c, sampled from a Gaussian distribution with mean 0.3 and standard deviation 0.1 (lower values clipped at 0). Distributions for synapses that become filopodia (grey) and spines (purple). The label *filopodia* is assigned to synapses with a mean (over 10 seconds) weight smaller than *w*_0_. Synapses with a value equal or higher than *w*_0_ are labeled as *spines*. **G2:** Distribution of weights after learning. **G3:** Scatter plot of final weight *w*_*i*_ as a function of *c*_*i*_.

To describe our model mathematically, we make use of *nonlinear temporally asymmetric* (nlta) STDP (Equation (3) in Methods, (Gü tig et al., 2003)). This rule generalizes both add- and mult-STDP via a single parameter *µ. µ* governs a phase transition such that below a critical value, *µ*_crit_, learning is effectively additive, while for *µ* > *µ*_crit_ learning is qualitatively similar to mult-STDP. This rule leads to a very natural implementation of our proposal: filopodia have small µ values, and thus experience additive learning, and spines have higher µ values, and therefore exhibit multiplicative dynamics. A way to do this is to make *µ* an increasing function of the synaptic efficacy (Methods, Equation (4)). Our specific choice is for *µ* to low-pass filter the synaptic strength w, such that it smooths out and temporally averages the synaptic efficacy. In addition, this particular implementation is consistent with the abovementioned experimental result (Vardalaki et al., 2022) that filopodia transition to spines if they keep a consistently high synaptic efficacy (as this leads to an increase in *µ*).

One of the benefits of using well-established models of learning is that one can make high-level predictions of the learning rule based on previous analyses of these models. Add-STDP is said to be *highly competitive*, while mult-STDP is instead *weakly competitive* (or *stable*) (Kempter et al., 1999; Song et al., 2000; Van Rossum et al., 2000; Gütig et al., 2003). These descriptions come from the concept of *competition* and *cooperation* that arise in mean-field models of STDP. Intuitively, competition refers to the fact that if one synapse increases its efficacy, it has a net depressive effect on all other synapses (Figure 1D, orange lines). This increase in depression is the same for all other synapses and is (with our stimulation protocol) independent of input correlations. Cooperation, on the other hand, has the opposite effect. When a synapse is potentiated, the average potentiation of the rest of the synapses increases. However, this increase in potentiation depends on the cross-correlations between presynaptic activity. In this sense, cooperation only favors potentiation of correlated synapses (Figure 1D, red lines). In both additive and multiplicative learning these two components exist, but with differences. In add-STDP competition and cooperation are weight-independent (Figure 1E, right subpanel). This leads to a so-called *strong competition*. In this scenario, learning splits synapses between those who *win* the competition, and accumulate at the upper hard bound, and those who *lose* it, and accumulate at zero. In mult-STDP, because the competition-cooperation ratio depends on the synaptic strength, weights are naturally allocated a continuous efficacy that reflects their level of correlation with the rest of the presynaptic pool. This results in a unimodal distribution that does not distinguish between *losers* and *winners*, and is thus called weakly competitive.

These notions of strong and weak competition can help predict the behaviour of FS-learning. Quantitatively, the terms competition and cooperation in STDP models refer to two mean-field terms that determine synapses’ evolution (Gü tig et al., 2003). We can therefore compute the competition and cooperation terms in FS-learning at the steady state (Methods, Equations (12) and (13)) and see how they change as a function of the synaptic efficacy w. If we compare the shape of the competition and cooperation in FS-learning with additive and multiplicative rules (Figure 1E small left and right subpanels, respectively), we observe the following: i) for weights between 0 and *w*_0_ (the arbitrary lower soft-bound of multiplicative learning), competition and cooperation are almost constant, resembling the curves of add-STDP (left subpanel), ii) when the weight crosses the *w*_0_ threshold the competition and cooperation terms are very similar to mult-STDP (right subpanel). Our interpretation is that FS-learning implements a two-stage (first strong, then weak) competition between synapses. Filopodia will experience an additive field, and only those correlated enough will win the competition, allowing them to cross the *w*_0_ threshold. If this value is maintained, their associated µ will converge to the multiplicative region, making these synapses spines. Because now they follow multiplicative dynamics, however, they will start competing weakly between them, thus creating a unimodal distribution correlation-dependent distribution, as does mult-STDP.

To test this mean-field prediction, we show a postsynaptic neuron following FS-learning in the presence of correlated input (Figures 1F and G). We use a conductance-based Leaky Integrate-and-Fire (LIF) neuron that contains 1000 excitatory and 200 inhibitory presynaptic inputs, each modelled as a Poisson process (Figure 1F). Excitatory connections are plastic and follow FS-learning (Equations (3), (4) in Methods). Presynaptic inputs are not independent from one another, but instead have a temporal correlation structure 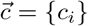 that determines how correlated the spike times of neuron *i* are to the rest of the presynaptic pool. To do this, we generate a reference spike train, and then *c*_*i*_ indicates how similar are the spike times of presynaptic neuron *i* is with that reference. Indirectly, this makes two neurons *i, j* that are very correlated with the reference spike train also very correlated between them, inducing cross-correlations of the order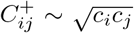 (Methods, Equation (19)). All presynaptic neurons have the same temporally averaged firing rate, denoted by *r*_pre_. As a proof of concept, we use here a correlation structure sampled from a Gaussian distribution (Figure 1G1). As predicted, due to the first stage of the competition, FS-learning results in two qualitatively distinct groups, filopodia (*w* < *w*_0_) and spines (*w* ≥*w*_0_). These two groups have very different averages compared to their variance (Figure 1G2). In addition, as also predicted by the second stage of the competition, weights *w*_*i*_ depend on the correlation value *c*_*i*_ of each synapse (Figure 1G3). Simulations support our two-stage competition proposal, which predicts an initial bimodal classification (first stage) and a continuous representation of the correlation strength in the synaptic efficacy (second stage).

### FS-learning represents input correlations better than add-STDP, and is more selective than mult-STDP

Having seen that FS-learning inherits properties of both additive and multiplicative learning, now we quantify how the receptive fields obtained via filopodia and spines compare to pure add- and mult-STDP. We choose von Mises shaped input correlations (Figure 2A) for our simulations for two reasons: First, the experimental study that motivates our research (Vardalaki et al., 2022) is based on visual cortex cells. Second, this choice gives input patterns a rich non-binary correlation structure, which allows us to test the correlation representational power. We stimulate our neuron for 200 seconds and then extract two measures: (i) the Pearson correlation *r* between the developed synaptic efficacies and the correlation strength of their corresponding presynaptic activity (Figure 2B and Methods), and (ii) the Discrimination Index *DI* that results from averaging the neuron response to different correlation patterns (Figure 2C and Methods). We use r as a proxy to how well the formed receptive fields (RFs) represent input correlations, and *DI* to quantify the ability of our neuron to discriminate different inputs.

**Figure 2.**
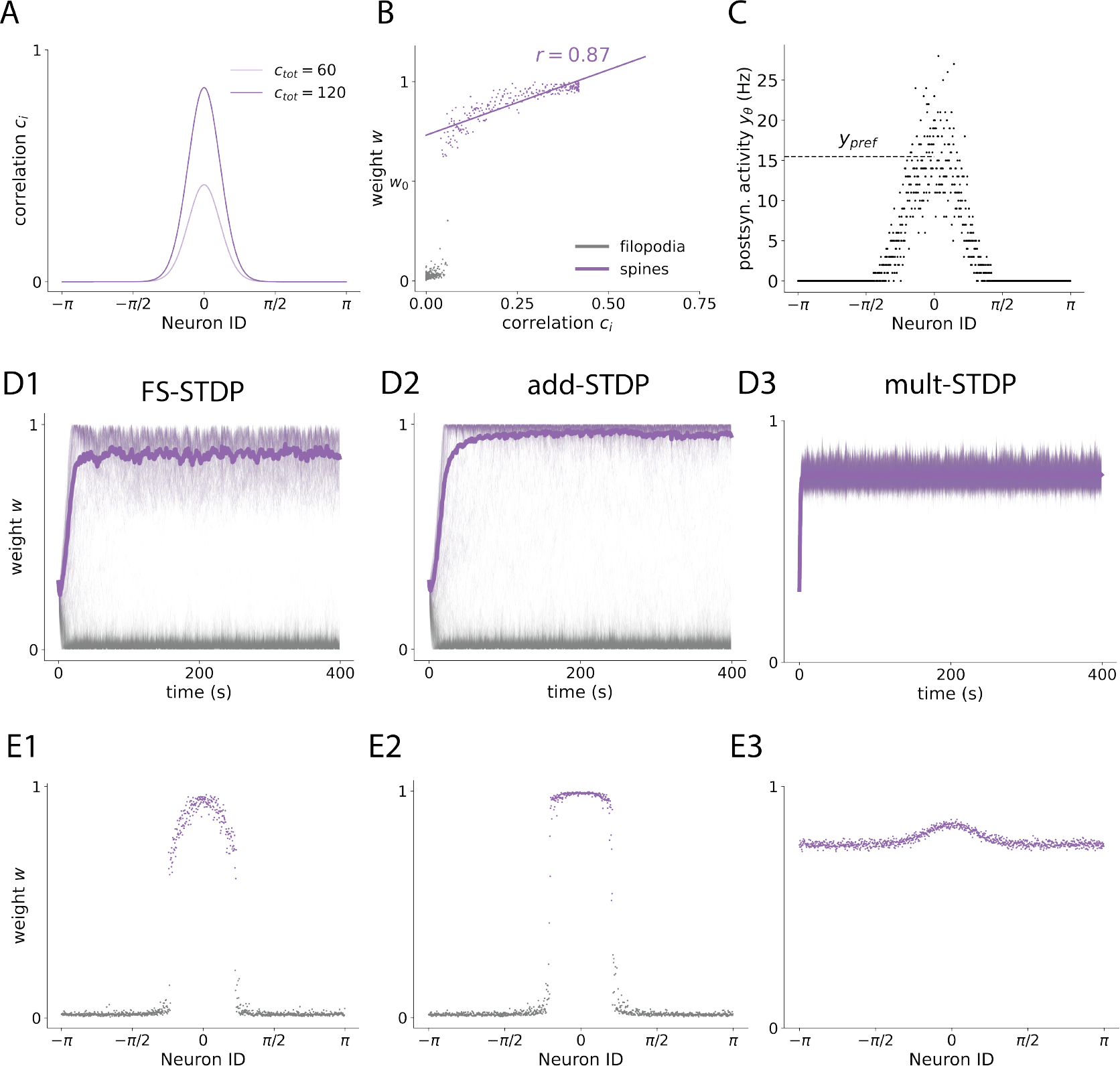
Receptive Fields (RFs) formed with FS-STDP inherit the bimodality and sparsity of add-STDP, and the correlation structure representation of mult-STDP. **A:** Input correlation structure used in our stimulation protocol. Each presynaptic neuron *i* is assigned an angle as a Neuron ID, and a correlation strength value *c*_*i*_ according to the pdf of a von Mises distribution at that angle. *c*_tot_ controls the total amount of correlation such that Σ *c*_*i*_ = *c*_tot_ (Methods, Equation (21)). **B:** Example scatter plot of weights after learning. To obtain r in Figure F we compute the Pearson correlation between spines’ synaptic efficacy and the corresponding *c*_*i*_ value. Grey corresponds to filopodia, and purple to spines. The straight line represents the linear regression of the synaptic efficacy after convergence of spines and the correlation strength *c*_*i*_. **C:** Example scatter plot of neuron output activity *y*_*θ*_ (sampled over 1 second) for different input correlation structures, after learning. Each point represents the postsynaptic firing rate when the input correlations are centered around a specific angle (the input pattern is a shift of the one in panel A such that the neuron with that ID has the highest *c*_*i*_). To reduce the bias in the discrimination index *DI*, the output activity at the preferred angle (Neuron ID = 0) is measured for 100 seconds. **D:** Example of weight trajectories for FS-STDP (**D1**), add-STDP (**D2**), and mult-STDP (**D3**). **E:** Example of receptive fields formed (weights at *t* = 400 s), same order as in **D**.

We start by noting the differences seen in both the weights trajectories and developed receptive fields (Figures 2D and 2E, respectively). Both FS and add learning result in very similar initial trajectories, which are associated with the first (strong) stage of the competition (2D1 and 2D2). In terms of the RFs formed, this initial strong competition makes both FS-STDP and add-STDP present a group of synapses with 0 efficacy along with a subgroup with significant synaptic strength (Figure 2E1 and 2E2). This is not the case of mult-STDP, where all of the synapses are located around a single mean (Figure 2E3). If one focuses on non-zero synapses, however, then FS-STDP is more similar to mult-STDP, as both learning rules continuously match the input correlation structure. In contrast, add-STDP erases this information and yields a binary RF resulting from classifying presynaptic input as correlated enough or not.

We quantify the differences in discriminability (as measured by DI) and input representational capacity (as measured by *r*) of FS-learning across different values of potentiation-depression imbalance (parameterized with α), and across different values of correlation strength (given by *c*_tot_). These two terms control the ratio between cooperation and competition, which strongly affects the formation the RFs (see Methods). We obtain *r* and *DI* for each combination of these two parameters, as well as for every learning rule (Figures 3A and 3B). *r* measures the strength of the linear relation between a synapse *w*_*i*_ and the correlation strength (*c*_*i*_) of its corresponding presynaptic neuron with the rest of the pool. Assuming no higher order relationships exist between the two, we take this metric to indicate how well the input correlations are imprinted into the weight structure, with *r* ranging from 0 (not well represented) to 1 (well represented). In general, there is a high degree of correlation in FS-learning (Figure 3A1). Due to the second stage of the competition, the input structure is much better imprinted in FS-STDP than in additive learning (Figure 3A2), reflected by higher *r* values for FS-STDP. In terms of discriminability, FS-STDP performs similarly to add-STDP (Figure 3B2, *FS* − *add*). The lack of synaptic specialization induces a discriminatory capacity of mult-STDP near zero.

**Figure 3.**
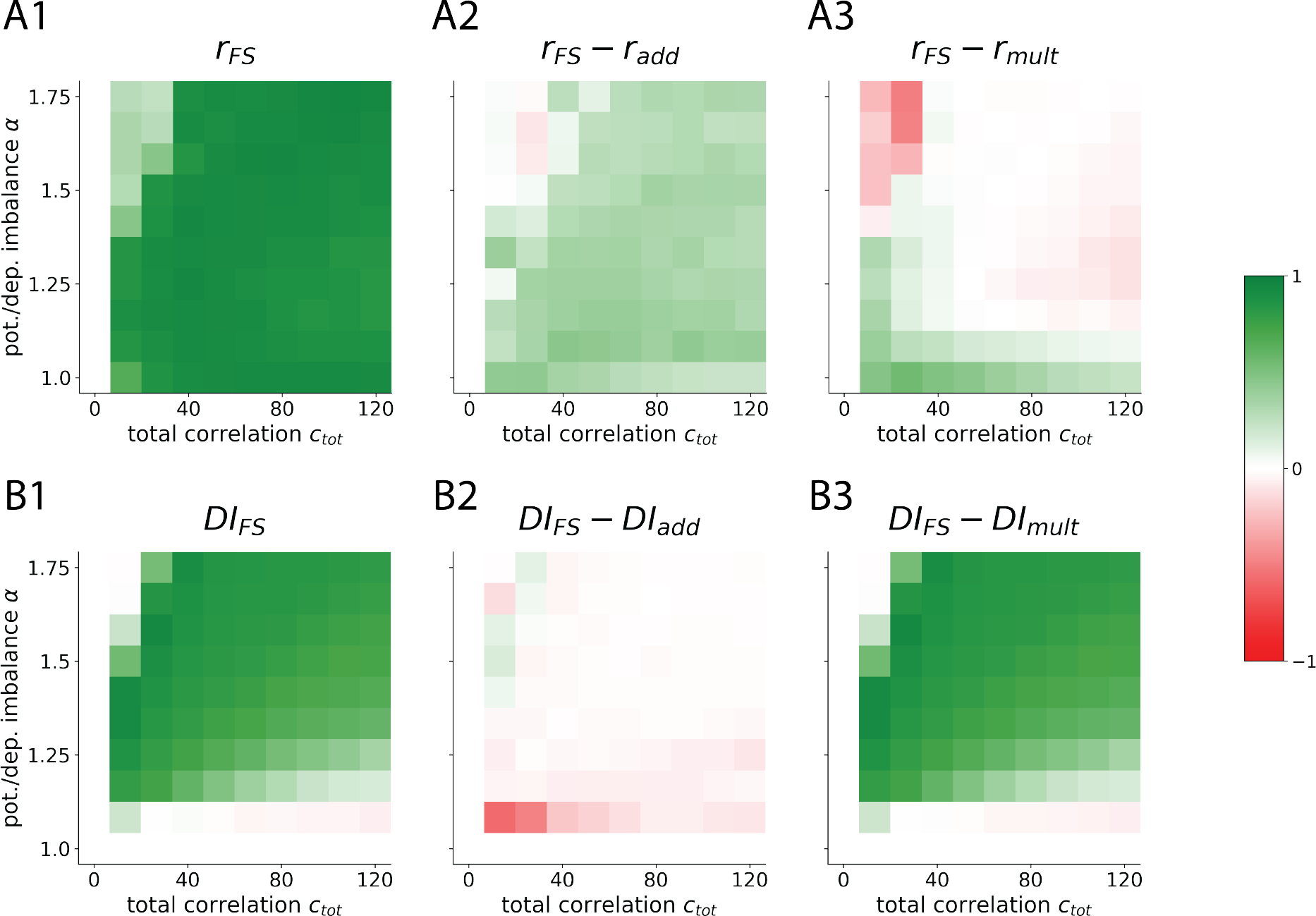
RF correlation representation (*r*) and discriminatory index (*DI*) across total correlation *c*_tot_ and potentation/depression imbalance *α* (FS-STDP and difference with add-STDP and mult-STDP). All values are averaged over 5 seeds. **A:** Heatmaps showing the Pearson correlation for *w*_*i*_ vs *c*_*i*_ (see Figure **B**). Subindex labels indicate what learning rule is being shown/compared (e.g. *r*_*F S*_− *r*_*add*_ means, *r* value for RFs obtained via FS-learning minus *r* value obtained via additive learning). x-axis shows increasing values of total correlation and y-axis increasing values of potentiation-depression imbalance *α*. Colormap ranges from Red to White to Green for values −1, 0, and 1. White regions found for small correlation values correspond to points in the parameter space where a RF has not (fully) formed. Red regions in panel F are points where a RF has formed with one rule but not the other. See Supplementary Figures 1, 2, and 3.**B:** Same for Discrimination Index averaged for all possible Neuron ID = *θ* (see Figure 2**C**).

Our results propose a mechanistic and biologically plausible explanation of how highly specialized RFs that continuously represent the input correlations can emerge via the strong competition of filopodia and the weak competition of spines. In addition, it explains how unimodal yet selective distributions could arise in the experimentally observed synaptic distributions (Barbour et al., 2007).

### FS-learning makes changes resistant to new correlated input

So far, we have restricted our simulations to the case where initial weights are all the same, and salient statistical patterns in the environment have not yet been imprinted in the weight structure. Usual environments, though, don’t have fixed statistics, and are subject to abrupt changes. This posits a trade-off whereby the brain needs to adapt to integrate new information, sometimes inconsistent with previous experience, but cannot just immediately through away any past memories. One classic solution to this problem is synaptic consolidation, which prevents memories from being erased after their formation (Fusi et al., 2005; Clopath et al., 2008; Barrett et al., 2009). In our model, synaptic consolidation is implemented with the existence of a lower soft-bound *w*_0_ (Equation (3) in Methods, also see Figure 1E), which makes depression approach zero when *w* → *w*_0_ from above. Whether this bound is effectively hard (impossible to trespass) or soft (difficult to trespass) is controlled by the *protection parameter a* (Methods, Equation (4)). Specifically, given a consistent synaptic efficacy *w*, greater *a* values will result in higher *µ* values at convergence. This will, in turn, result in a *harder* soft-bound, and increase the protection of the receptive field.

To investigate how changes in input correlations affect the preexisting synaptic structure we apply a new stimulation protocol. In this scenario, there exist two non-overlapping groups of input patterns, A and B (purple and yellow in Figure 4A, respectively). Pattern A and B are orthogonal in the sense that the neurons with nonzero correlation are disjoint. We start with pattern A for 200 seconds, and at then we change the correlation structure from pattern A to pattern B for 400 seconds more. Applying this protocol, one can encounter three different scenarios: (i) the previous memory is erased with the new pattern taking over (Figure 4B2, *Total Overwriting*), (ii) the previous memory is maintained, but pattern B is also imprinted (Figure 4B3, *Partial Overwriting*), and (iii) the previous memory is maintained, and the new pattern is not imprinted (Figure 4B4, *No Overwriting*). We classify the trajectories obtained for each simulation as one of these three qualitatively different cases. To do that, we compare the RFs formed with pattern A or B alone 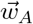 and 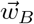 respectively) with the one formed when first A and then B is presented 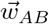 (Figure 4C and Equation (24) in Methods).

**Figure 4.**
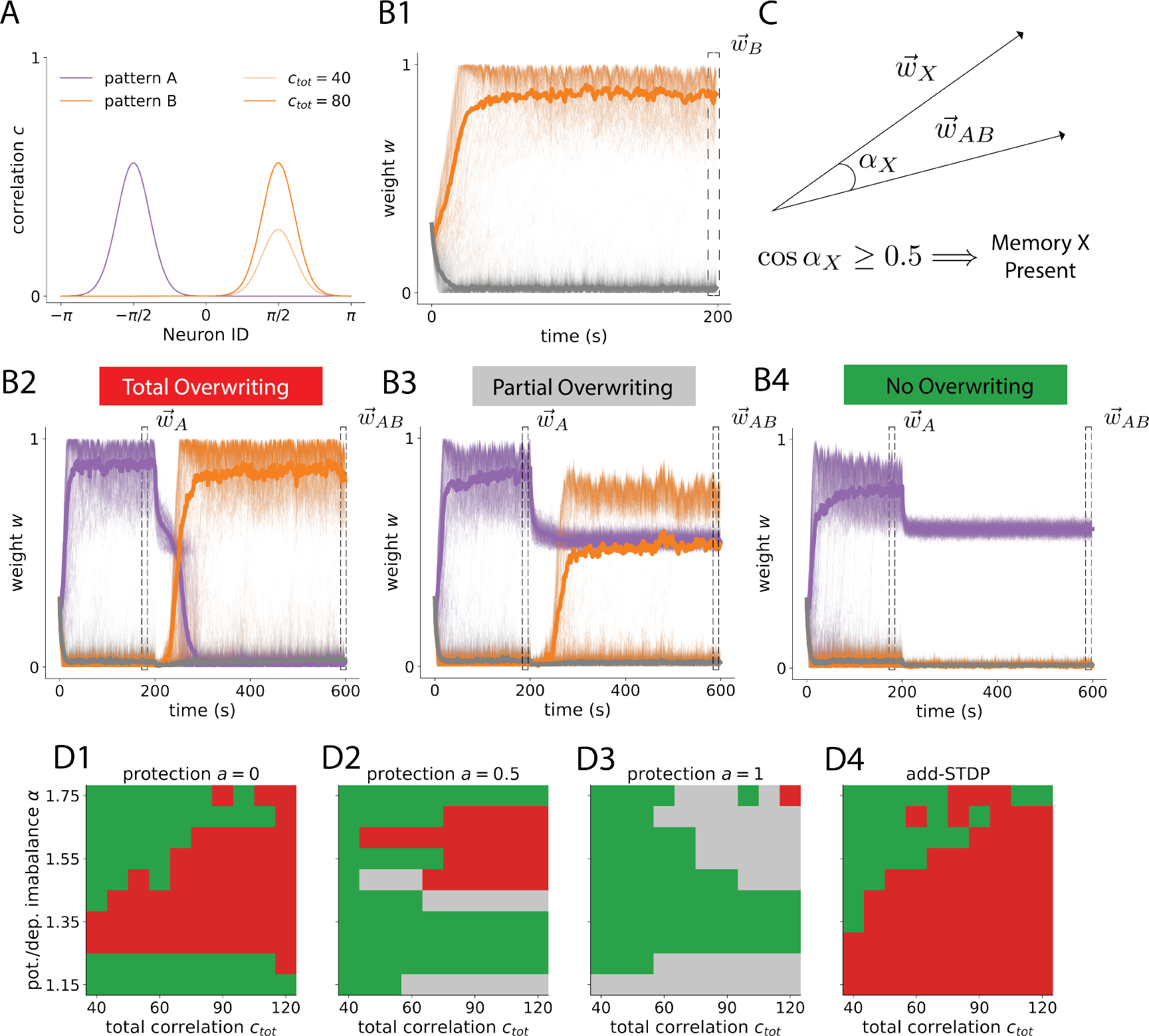
**A:** Stimulation protocol. Input neurons have an input correlation profile as in purple (pattern A) for the first 200 seconds of the simulation. Then, it switches to the yellow correlation profile (pattern B, shifted by *π* with respect to pattern A). **B:** There exist different scenarios: (i) The new memory (which would lead to **B1** in the absence of previous structure) takes over the previous one (red, **B3**). (ii) The new memory is imprinted but the old RF is conserved (orange, **B4**). (iii) The original RF is preserved and the new memory is not imprinted (green, **B5**). Orange traces correspond to weights that become spines with pattern B alone, and purple ones weights that become spines with pattern A alone. **C:** By comparing the final RF 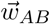 with that corresponding to pattern X alone (X = A,B), one can obtain a similarity score that determines whether the memory X is present or not (Methods, Equation (24)). **D:** Classification of receptive fields formed across potentiation/depression imbalance α and total correlation *c*_*tot*_ for different protection parameter a values in FS-STDP (**D1, D2**, and **D3**) and in add-STDP (**D4**). Colors as indicated in panels **B2, B3, B4**. Ranges of *α* and *c*_*tot*_ were chosen so that a RF was always formed (see Figure 3 and Supplementary Figures 1 and 2).

We study how the above-mentioned regimes are affected by the total amount of correlation of pattern B *c*_*tot*_, the potentiation/depression imbalance *α*, and the protection parameter *a*. Higher levels of correlation in pattern B facilitate the formation of the new pattern, as the new correlated synapses can cooperate more strongly. This leads to an abundance of transitions from no overwriting to partial or total overwriting with *c*_*tot*_ given a fixed α (Figure 4D). The effect of increasing the imbalance between potentiation and depression is not trivial, as increasing depression prevents the new memory to take over but also increases the pressure over the previously formed memory. In consequence, and as shown in simulations, increasing *α* can drift the system from memory protection to memory overtake but also the opposite, with the process being also coupled to the specific amount of correlation of pattern B (Figure 4D). Increasing protection parameter a consistently prevents the previously formed memory (more extended green or blue regions for higher values of a, Figure 4D). Nevertheless, doing so also decreases the competition between filopodia and spines, thus favoring the formation of the new RF while the previous one is maintained (changes from green to orange). Compared to additive STDP, FS learning has two advantages: first, it allows a non-existing regime of memory linking/overlapping (Figure 4D4). Secondly, it allows transitioning from an add-like scenario (Figures 4D1 and 4D4) to regimes in which previously formed memories are more and more protected (Figures 4D2 and 4D3).

Altogether, this shows a richness of possible interactions between previous and new environmental statistics in FS-STDP. Our learning rule allows for an intermediate state in which receptive fields aggregate (orange), not allowed in classic models of STDP. Furthermore, it provides an active protection mechanism that goes beyond quiescence and preserves weight structure in the presence of postsynaptic activity. Interestingly, these different regimes can be controlled with model parameters like *α* or a, so one could imagine these being physiologically driven to favor one or another depending on developmental stages or the measured environmental uncertainty.

## Discussion

We have presented a computational model that describes how filopodia and spines are differently affected by plasticity, as well as how they transition from one to the other. Our learning rule, Filopodium-Spine STDP, posits that highly volatile and plastic filopodia exhibit additive-like plasticity, whereas spines follow multiplicative learning dynamics. We use nonlinear-temporally-asymetric (nlta) learning to generalize the learning rules that follow both silent synapses (filopodia) and spines, with the addition of weight dependence in the parameter that governs the additiveness and multiplicativeness. FS-STDP contains two key ingredients that make it functionally appealing: (i) it is able to selectively encode in the synaptic state a rectified version of the correlation structure of input and (ii) it acts as a memory consolidation mechanism by protecting previously formed receptive fields.

We have shown that the encoding properties of our learning rule come from a two-stage competition (strong, followed by weak), such that synapses are first biomdally distributed (as in add-STDP) and then also continuously represent the correlation structure of the input. This comes as an alternative solution to the *Stability vs Neuronal Specialization* dilemma (Gilson & Fukai, 2011). Here we have focused on weights between zero and one, and have excluded the possibility of further representing the correlation structure using long-tail distributions. A similar two-stage learning could be implemented by imposing a parameterized log dependence such that filopodia follow add-STDP and spines log-STDP. Our model, together with experiments, is in accord with the hypotheses that optimal network capacity can be obtained when a large fraction of the synapses is *silent* (Brunel et al., 2004) and the experimental sparseness of network connectivity (Song et al., 2005). Furthermore, FS-STDP proposes a mechanistic explanation of how this sparse connectivity might be obtained in the first place, and how it can flexibly adapt to changes in the environment statistics. In a sense, this effectively makes the first stage of FS learning a *structural plasticity rule* (Knoblauch et al., 2014; Ozcan, 2017).

The second property of FS-learning is the protection of formed receptive fields in the presence of new input correlations. This fits in the literature of synaptic consolidation, as it can be understood as a cascade model (Fusi et al., 2005) where the *spineness* (controlled by parameter µ) is an internal variable of the weight. It can also be placed in the more recent context of bidirectional dynamic processes (Benna & Fusi, 2016), where the weight pushes-pulls µ and in turn high values µ stabilize the weight. We have restricted our analysis to the interplay between fliopodia and spines, but the model could be extended to include other well-known processes of synaptic consolidation that would affect spines (as could be the experimentally (Frey & Morris, 1997; Redondo & Morris, 2011) and computationally (Clopath et al., 2008; Ziegler et al., 2015; Lehr et al., 2022) tested Synaptic Tagging and Capture (STC) mechanism).

FS-learning can also account for spine volatility (Mongillo et al., 2017). If the µ value associated with the lower soft-bound *w*_0_ leads to additive dynamics, then not all spines are protected. Instead, they decay even in the absence of new correlated input. This would happen in order of previous correlation strength (less correlated decay first) until the competitive term is weak enough. This would imply a synaptic turnover rather active (instead of passive) making less relevant spines decay into filopodia first.

While we do not examine directly the implementation of FS-learning in a recurrently connected network, we hypothesize that it could be beneficial for learning neuronal assemblies. While the advantages of graded synapses in memory formation via attractors have been described before (Satel et al., 2009), the representational power of FS-learning could also lead to imprinting more complex conceptual structures. Assembly formation in recurrent networks is usually limited to independent groups of neurons that represent orthogonal input, as classic models of competitive learning would otherwise lead to representational collapse and merging of assemblies. FS-STDP could potentially overcome this by distinguishing between-assemblies and inter-assemblies synaptic strengths, while maintaining a sparse connectivity. The intrinsic protection that our model gives to spines could additionally make the assemblies formed resistant to new correlated inputs that would otherwise erase them (preventing catastrophic forgetting).

Our model assumes that the same process of filopodia being converted into spines when they are potentiated also exists in reverse (while only the first has been experimentally observed). Future experiments where this inverse transformation is investigated. However, our conversion of spines into filopodia could still be considered a simplification of a renewal process in which a spine disappears and randomly a close-by filopodium is created. This study also naturally suggests a systematic characterization of spike-timing-dependent plasticity in filopodia, which could help confirm if they actually follow additive dynamics and how the learning kernels vary across the filopodium-spine spectrum. We have also not modelled AMPA and NMDA channels separately, while filopodia are known to contain NMDA channels even in the absence of AMPA receptors (Jontes & Smith, 2000; Vardalaki et al., 2022). We speculate that including this distinction could be beneficial for the formation of neuronal assemblies in recurrent networks, given the slow dynamics of these channels and their role as coincidence detectors (Seeburg et al., 1995; Tabone & Ramaswami, 2012).

In summary, we have presented a model of synaptic plasticity that distinguishes between two synaptic structures: filopodia and spines. We have assumed that filopodia follow highly competitive additive STDP and spines learn according to multiplicative STDP, which leads to the formation of sparse but graded receptive fields that continuously represent input correlations. Furthermore, our model proposes that the transformation of filopodia into spines can be seen as a first element of synaptic consolidation, thus making the synaptic weights learned protected from changes in environment statistics.

## Methods

### Neuron Model

We use a conductance-based Leaky Integrate-and-Fire model with excitatory and inhibitory input. Passive membrane potential dynamics are described by the following equation:

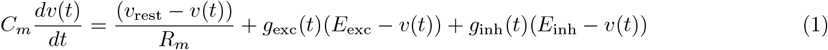

where the conductance response to presynaptic spikes is shaped using alpha functions:

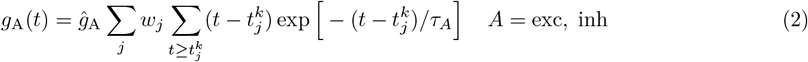

If at time 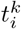 neuron *i* has membrane potential *v* = *v*_th_, then *v* is reset to *v*_rest_ at 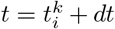 and 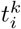 included in the spike train *S*_*i*_(*t*). Parameter descriptions and values used in simulations are specified in Table 1.

**Table 1:** Neuron Parameters.

| Symbol | Value | Description |
| --- | --- | --- |
| $v$ | Variable (mV) | Membrane Potential |
| $C_m$ | 200 pF | Membrane Capacitance |
| $R_m$ | 100 MΩ | Membrane Resistance |
| $v_{\text{rest}}$ | -70 mV | Resting Potential |
| $E_{\text{exc}}$ | 0 mV | Excitatory Reversal Potential |
| $E_{\text{inh}}$ | -70 mV | Inhibitory Reversal Potential |
| $\hat{g}_{\text{exc}}$ | 0.15 nS | Excitatory Conductance Amplitude |
| $\hat{g}_{\text{inh}}$ | 0.25 | Inhibitory Conductance Amplitude |
| $\tau_{\text{exc}}$ | 5 ms | Excitatory Time Constant |
| $\tau_{\text{inh}}$ | 5 ms | Inhibitory Time Constant |

### Synaptic Plasticity Model

#### Filopodium-Spine STDP

Our model adds intrinsic dynamics to the parameter *µ* introduced in Gü tig et al. (2003), making it coupled to the weight *w*_*i*_ (i denotes presynaptic index, there is only one postsynaptic neuron):

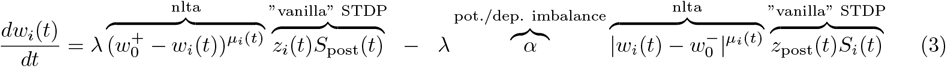

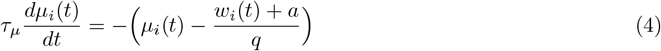

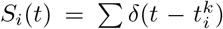 are the spike trains of presynaptic neuron *i* and *S*_post_ the spike trains of the only postsynaptic neuron. The pre (sub-index *i*) and post (sub-index post) synaptic traces are low-pass filtered versions of the corresponding spike trains:

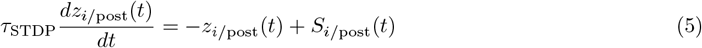

These equations combined implement a special weight-dependence such that only smalls weights experience strong competition. In Gütig et al. (2003), it is shown that, given a fixed *µ*, learning can be more add-like (strongly competitive, if *µ* is below a critical value *µ*_crit_) or more mult-like (weakly competitive, if *µ* is above *µ*_crit_). Parameters *q* (quotient) and *a* (protection) are chosen such that learning goes from additive to multiplicative when weights go from 0 to approximately or higher than *w*_0_. Throughout this study, we let 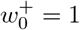 and simply write 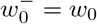 All parameters used in simulations are specified in Table 2.

**Table 2:**
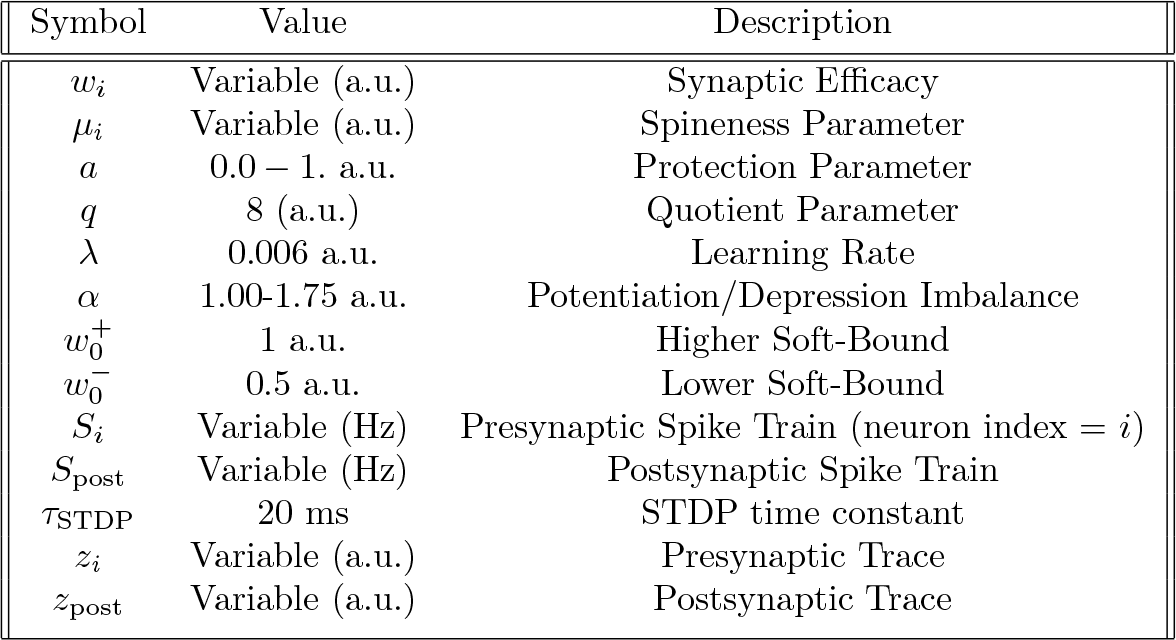
Synaptic Plasticity Parameters.

#### Additive STDP

We define additive STDP (add-STDP) as in Song et al. (2000):

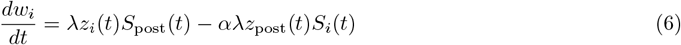

using same parameter values as in Table 2.

#### Multiplicative STDP

We define multiplicative STDP (mult-STDP) as in Van Rossum et al. (2000):

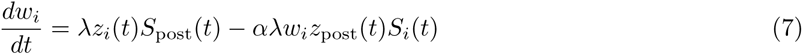

using same parameter values as in Table 2.

### Mean synaptic dynamics in a linear Poisson neuron

Following Gütig et al. (2003), we recover some of the main results assuming a Poisson linear neuron, which allows for an exact solution of the weight mean-field dynamics. In this context, the output neuron is the realization of an inhomogeneous Poisson process with an instantaneous firing rate

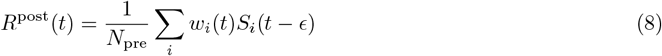

where *N*_pre_ is the number of presynaptic neurons and *ϵ* a small constant delay. Under these conditions, and if one has a stimulation protocol without backward correlations (as is our case, see below), one can obtain the mean-field description of the time evolution of *w*_*i*_:

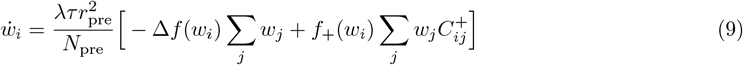

where *r*_pre_ is the presynaptic firing rate, *f*_+_(*w*_*i*_) and *f*_*−*_(*w*_*i*_) are the time-difference-independent amplitudes of potentiation and depression (respectively) and 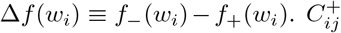 and is called the *integrated normalized cross-correlation* and is a function of the *normalized cross-correlation* 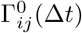 and are defined as:

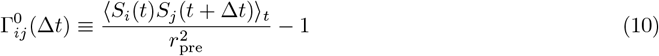

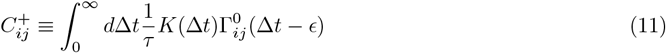

with *K*(∆*t*) the *learning kernel*, which defines how the weight changes as a function of the spike-time difference between pre and postsynaptic spikes (Figure 1B).

### Competition and cooperation in FS-learning

Equation (9) gives an intuition of the effect of increasing a synaptic efficacy *w*_*k*_. The negative term will become bigger for all synapses *w*_*i*_ due to the increase in Σ*w*_*j*_ (i.e. competition). On the other hand, the positive term will become higher, but scaled by the cross-correlations *C*_*ik*_ that exist between neurons *i* and *k* (cooperation). Thus, increases in synaptic efficacy will have a net negative or positive effect on other synapses depending on the correlation structure. By combining Equations (3) and (4) at convergence 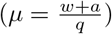 with Equation (9), one gets:

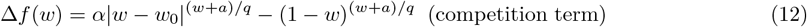

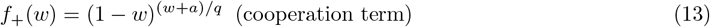

being ∆*f*(*w*) the competition term, and *f*_+_(*w*) the cooperation term. Note how competition is directly influenced by the imbalance between potentiation and depression *α*.

### Generating temporally correlated spike trains

To correlate a pool of presynaptic Poisson neurons, we generate a reference spike train:

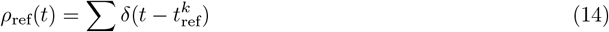

via a homogeneous Poisson process: 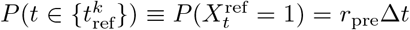 (∆t a small timestep). Now, for each neuron *i* in with correlation strength *c*_*i*_, the probability of firing at time *t* depends on whether that time is included in the reference spike train or not:

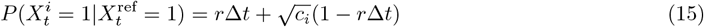

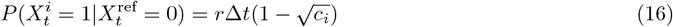

This ensures an average firing rate *r* over the defined period of time, while also imposing the following instantaneous cross-correlations:

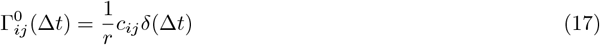

where *c*_*ij*_ denotes the pair-wise correlations

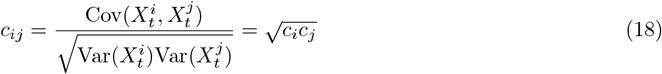

In turn, this results in the following integrated normalized cross-correlations, which define the level of cooperation between synapses:

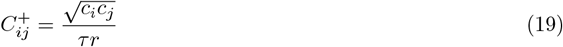

Note how here we have generalized to an arbitrary correlation structure 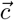, so not all neurons within a correlated pool are necessarily equally correlated.

### von Mises shaped correlation

The von Mises distribution is often used to describe tuning to different inputs that have a circular or periodic relationship. Its density function over an angle *θ* is given by

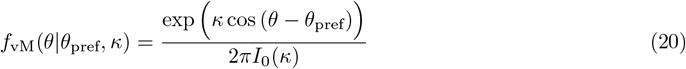

with *I*_0_(*κ*) = exp (*κ* cos *θ*)*dθ. θ*_pref_ denotes the center of the distribution (of preferred angle) and *κ* controls its width (or variance).

Given a Neuron ID = *θ*_*i*_, we assign each presynaptic neuron a correlation strength *c*_*i*_ = *f*_vM_(*θ*_*i*_) and then normalize according to parameter *c*_tot_ such that:

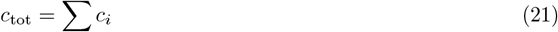

Parameter *c*_tot_ gives a measure of the total drive that the postsynaptic neuron effectively receives due to presynaptic temporal correlations.

### Discrimination Index

We define *y*_*θ*_ as the firing rate of the output neuron in the presence of an input correlation structure centered at *θ*. Then, the *Discrimination Index* at angle *θ* (*DI*(*θ*)), of a neuron trained with input centered at *θ* = pref, is

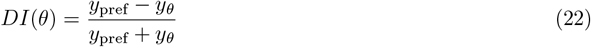

We then define the general Discrimination Index (*DI*) as:

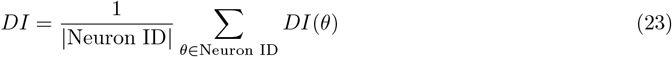

That is, the discrimination index evaluated at *θ* averaged over all possible values of *θ*.

### Memory Overlap

Given a set of conditions *X*, and a set of conditions *Y*, the *memory* after convergence is defined as 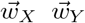 (respectively). Then, we measure the overlap between both memory via their cosine similarity:

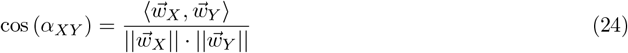

where 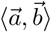 is the dot product of vectors 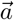 and 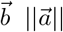 is the norm 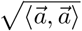 of 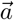, and *α*_*XY*_ is the angle between 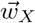 and 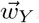 We use a threshold of 0.5 to determine whether a memory is present or not. The threshold was chosen such that the colormap in Figure 4D qualitatively corresponds with the regimes presented in Figures 4B2, B3 and B4 (total, partial or no overwriting). Also see Supplementary Figures 4, 5, 6, and 7.

### Implementation Details

We use BRIAN2 (Stimberg et al., 2019) in our simulations. Additional implementation parameters used in simulations are specified in Table 3.

**Table 3:** Simulation Parameters.

| Symbol | Value | Description |
| --- | --- | --- |
| $\Delta t$ (int.) | 0.5 ms | Integration timestep |
| $\Delta t$ (input) | 1 second | Input timed array width |
| $\Delta t$ ($w$ recording) | 1 second | Weight recording timestep |
| $w$ (initial) | 0.3 a.u. | Initial synaptic efficacy |
| $\kappa$ | 8 a.u. | von Mises pdf width parameter |
| $r_{\text{pre}}$ (default) | 35 Hz | Presynaptic firing rate (training) |
| $r_{\text{pre}}$ (discrimination index) | 10 Hz | Presynaptic firing rate (testing) |
| $r_{\text{pre}}$ (inhibition) | 10 Hz | Presynaptic firing rate (inhibitory neurons) |
| Figure | Variable | Value |
| 1 - B | $a$ | 0.5 |
| 1 - E | $a$ | 0, 0.5, 1 (thinner to thicker) |
| 1 - GX | $a$ | 0 |
| 1 - GX | $\alpha$ | 1.35 |
| 2 | $a$ | 0 |
| 2 | $\alpha$ | 1.35 |
| 2 - B, C, DX, EX | $c_{\text{tot}}$ | 60 |
| 3 - AX, BX | $a$ | 0 |
| 4 - BX | $\alpha$ | 1.35 |
| 4 - BX | $c_{\text{tot}}$ (pattern A and pattern B) | 80 |
| 4 - B1, B2 | $a$ | 0 |
| 4 - B3 | $a$ | 0.4 |
| 4 - B4 | $a$ | 1 |

### Presynaptic firing rates

Inhibitory neurons have a fixed 10 Hz firing rate. For excitatory neurons, we use a presynaptic firing rate (*r*_pre_) of 30 Hz in all our simulations except for computing the Discrimination Index (*DI*). A relatively high (30 Hz) presynaptic rate speeds-up simulations and avoids quiescence modes with an absence of plasticity. For example, in the simulations of Figure 4, a low presynaptic rate would lead to an effective *No Overwriting* regime because the postsynaptic neuron is silent and the weights are frozen. However, we are interested in the case where the learning rule inherently protects the RF even with ongoing plasticity. At high presynaptic firing rates, however, the postsynaptic neuron firing rate does not depend on the selectivity of the RF to the presented pattern. This is because the neurons with non-zero weight are able to excite the postsynaptic neuron even in the absence of correlations. To use the *DI* as a selectivity measure, which has the advantage of being normalized, we use a presynpatic firing rate of 10 Hz once the RF has formed.

### Use of total correlation *c*_tot_ and potentiation/depression imbalance α as parameter space

If one takes the mean-field learning dynamics (Equation (9)) and substitutes the cross-corelations 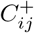 obtained in Equation (19), the following expression is obtained:

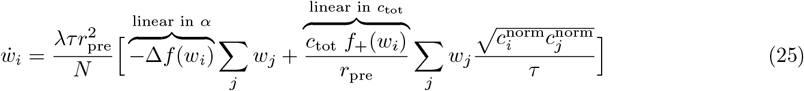

where 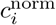are the normalized correlation values 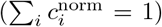 which are fixed given *θ*_pref_ and *κ*. One can see that the effect of increasing *r*_pre_ is equivalent to: (i) quadratically increasing the learning rate and (ii) inversely decreasing cooperation. Instead of changing *r*_pre_ (which would have with the aforementioned inconveniences), we sweep over the parameters *α* and *c*_*tot*_. This allows exploring different ratios of competition (modulated by α) and cooperation (modulated by *c*_tot_), while yielding similar stability and training times.

## Code Availability

The code for reproduction of the figures in the paper can be found online at https://github.com/albesagonzalez/filopodium-spine-learning

## Acknowledgments

This work was supported by BBSRC BB/N013956/1, BB/N019008/1, Wellcome Trust 200790/Z/16/Z, Simons Foundation 564408 and EPSRC EP/R035806/1.

## Supplementary Figures

**Figure 1:**
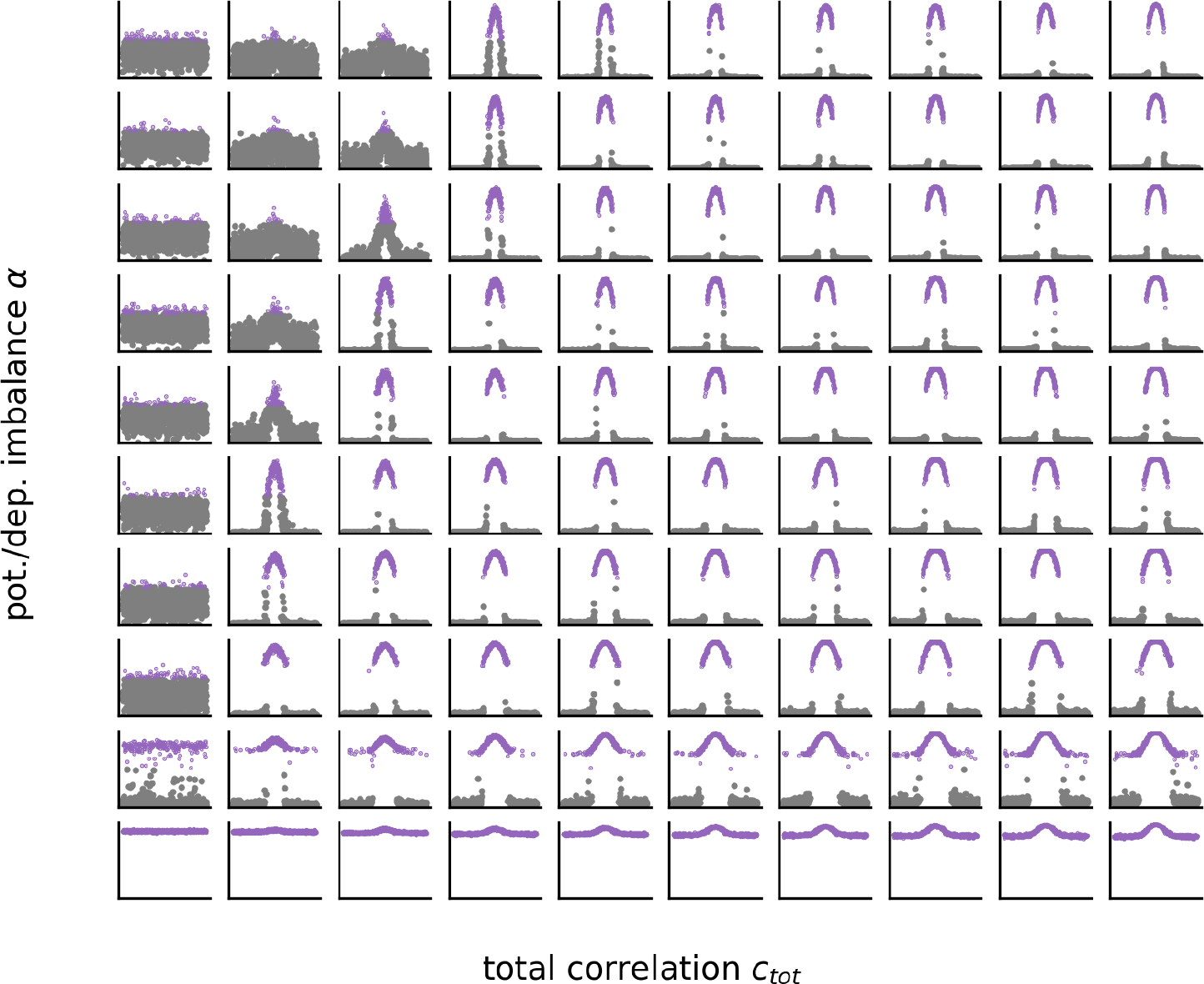
Receptive Fields after convergence for FS-STDP. Each subpanel corresponds to each of the 10×10 pixels in the heatmaps shown in Figures 3A1 and 3B1.

**Figure 2:**
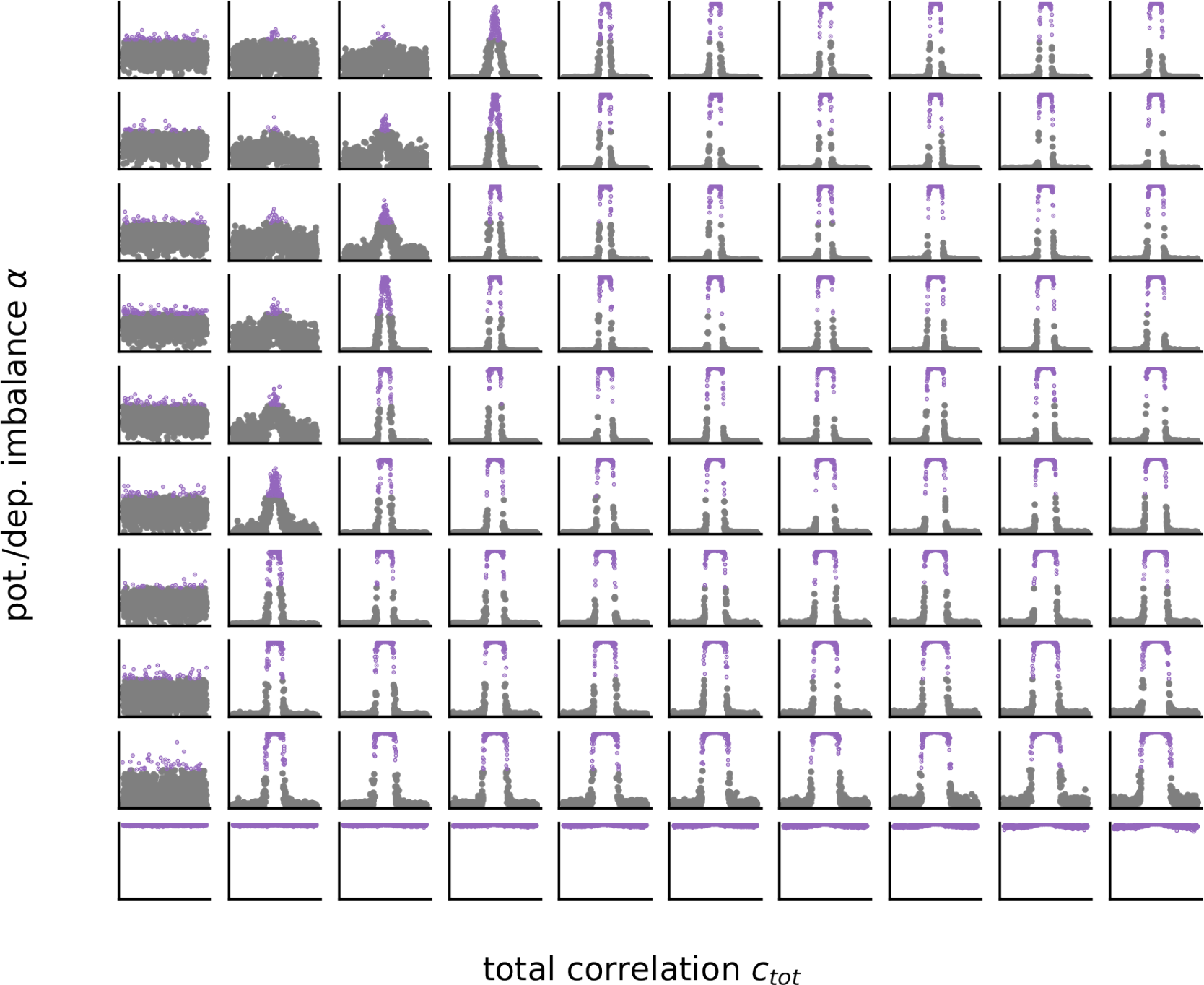
Receptive Fields after convergence for add-STDP. Each subpanel corresponds to each of the 10×10 pixels in the heatmaps shown in Figures 3A2 and 3B2.

**Figure 3:**
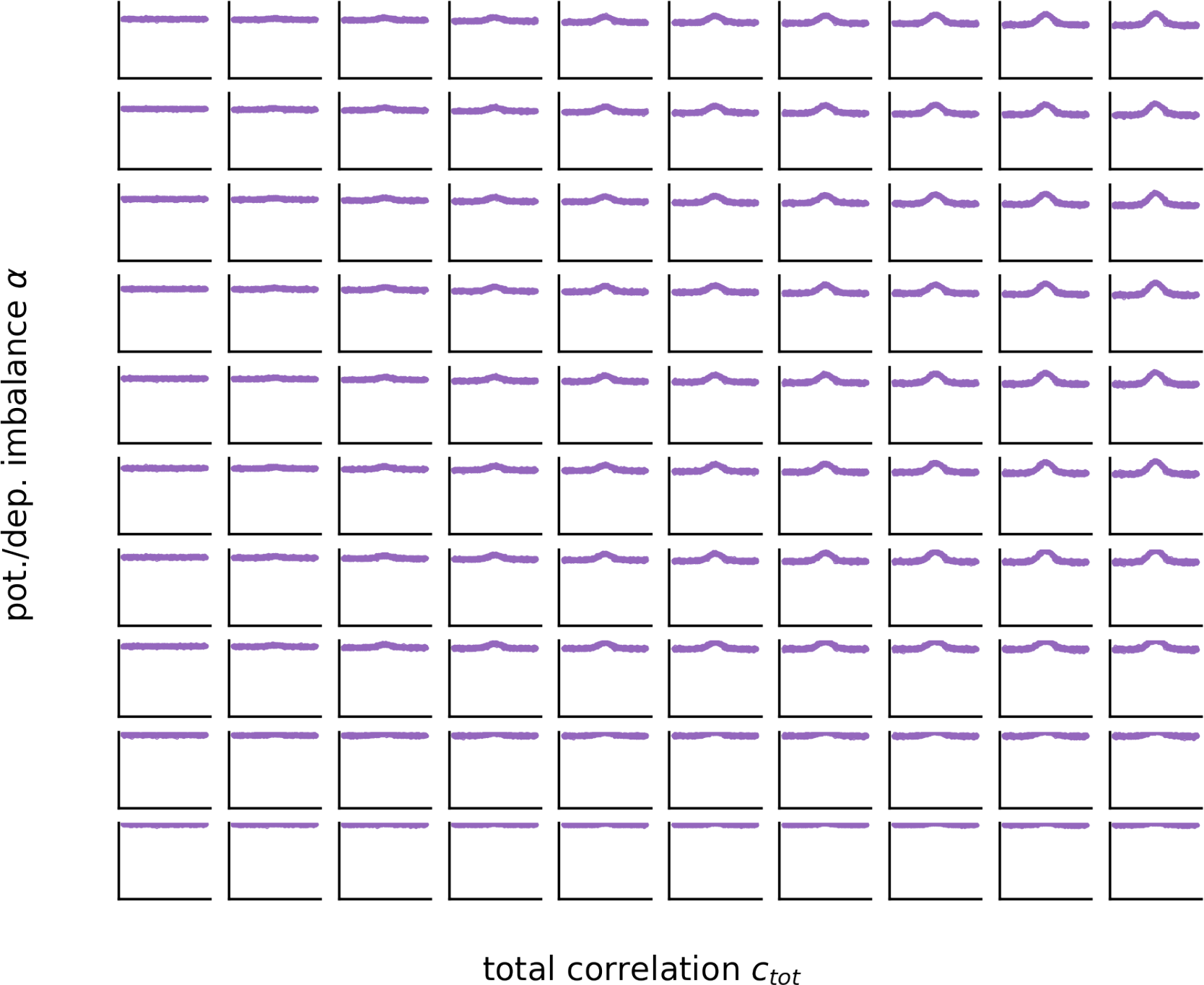
Receptive Fields after convergence for mult-STDP. Each subpanel corresponds to each of the 10×10 pixels in the heatmaps shown in Figures 3A3 and 3B3.

**Figure 4:**
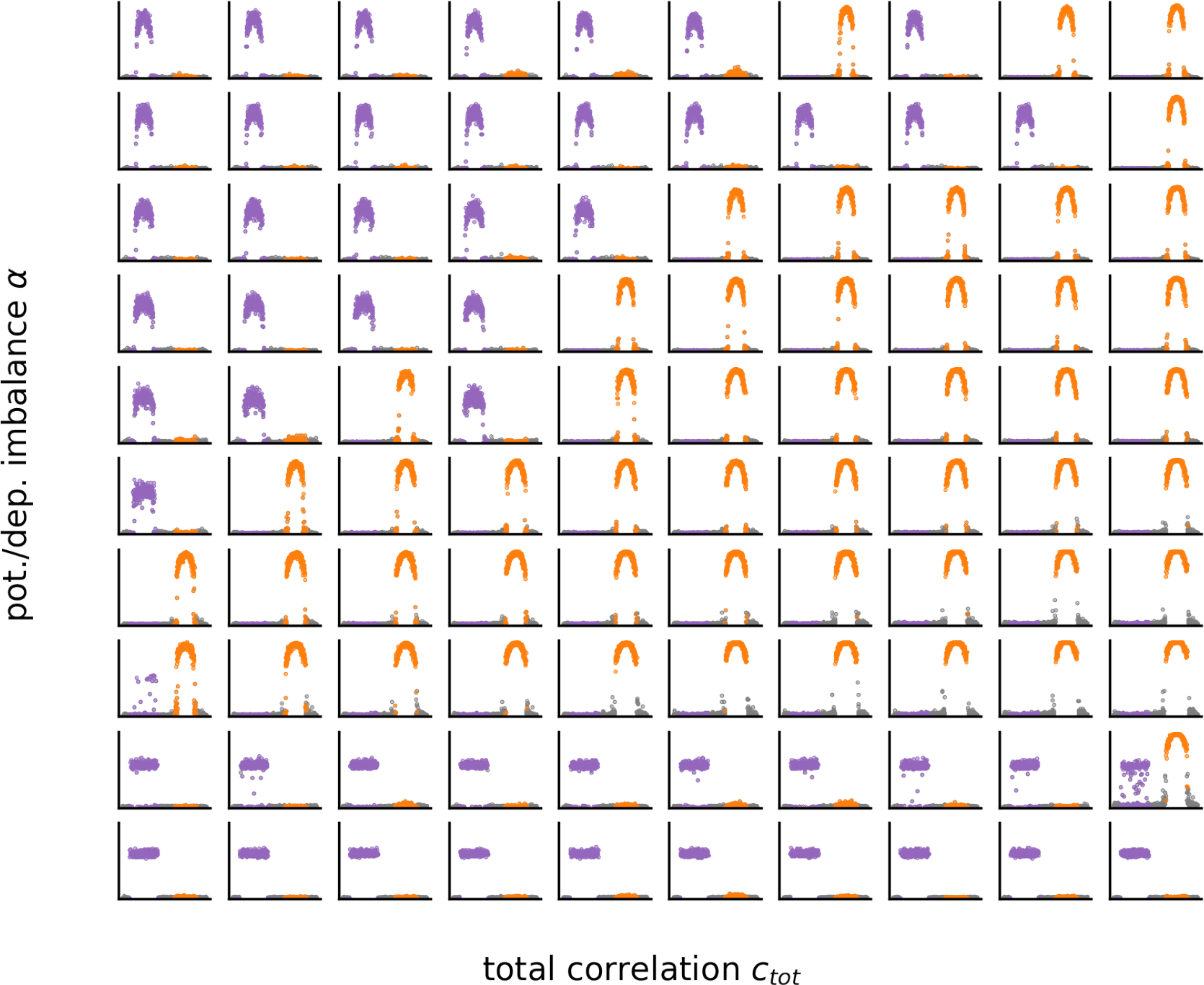
Receptive Fields after convergence for FS-STDP, a = 0 (pattern A and then pattern B). Each subpanel corresponds to each of the 10×10 pixels in the heatmaps shown in Figure 4C1.

**Figure 5:**
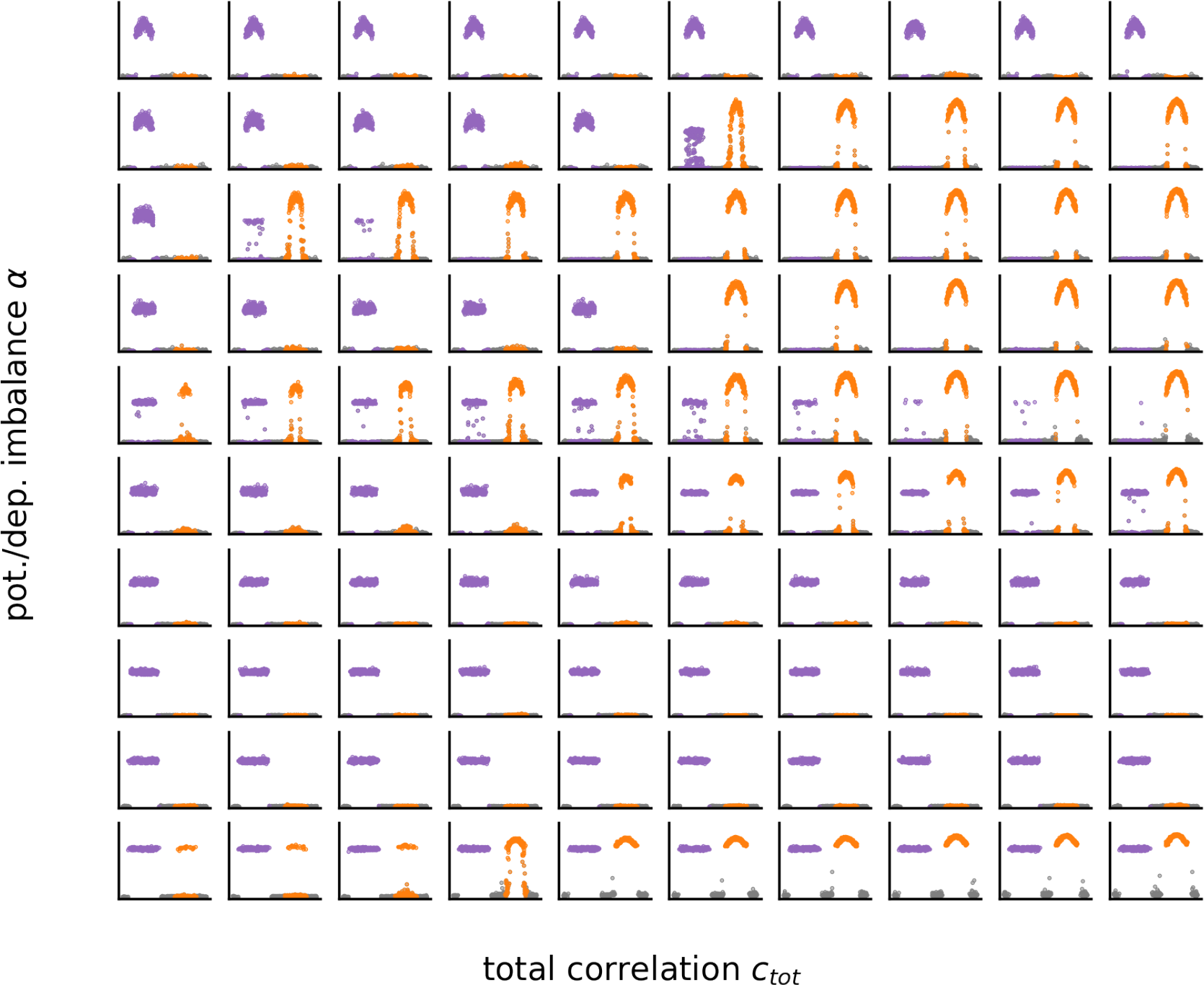
Receptive Fields after convergence for FS-STDP, a = 0.5 (pattern A and then pattern B). Each subpanel corresponds to each of the 10×10 pixels in the heatmaps shown in Figure 4C2.

**Figure 6:**
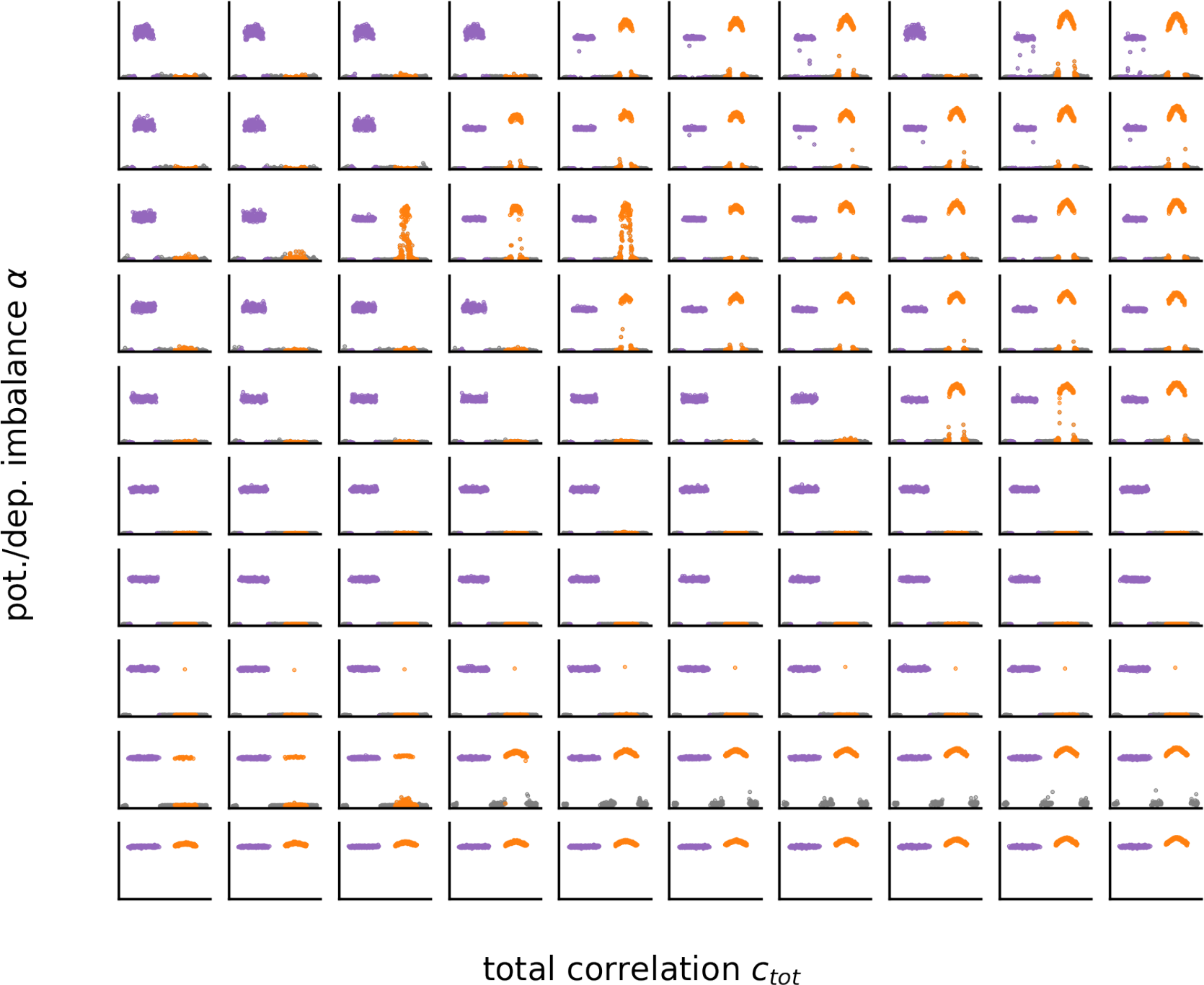
Receptive Fields after convergence for FS-STDP, a = 1 (pattern A and then pattern B). Each subpanel corresponds to each of the 10×10 pixels in the heatmaps shown in Figure 4C3.

**Figure 7:**
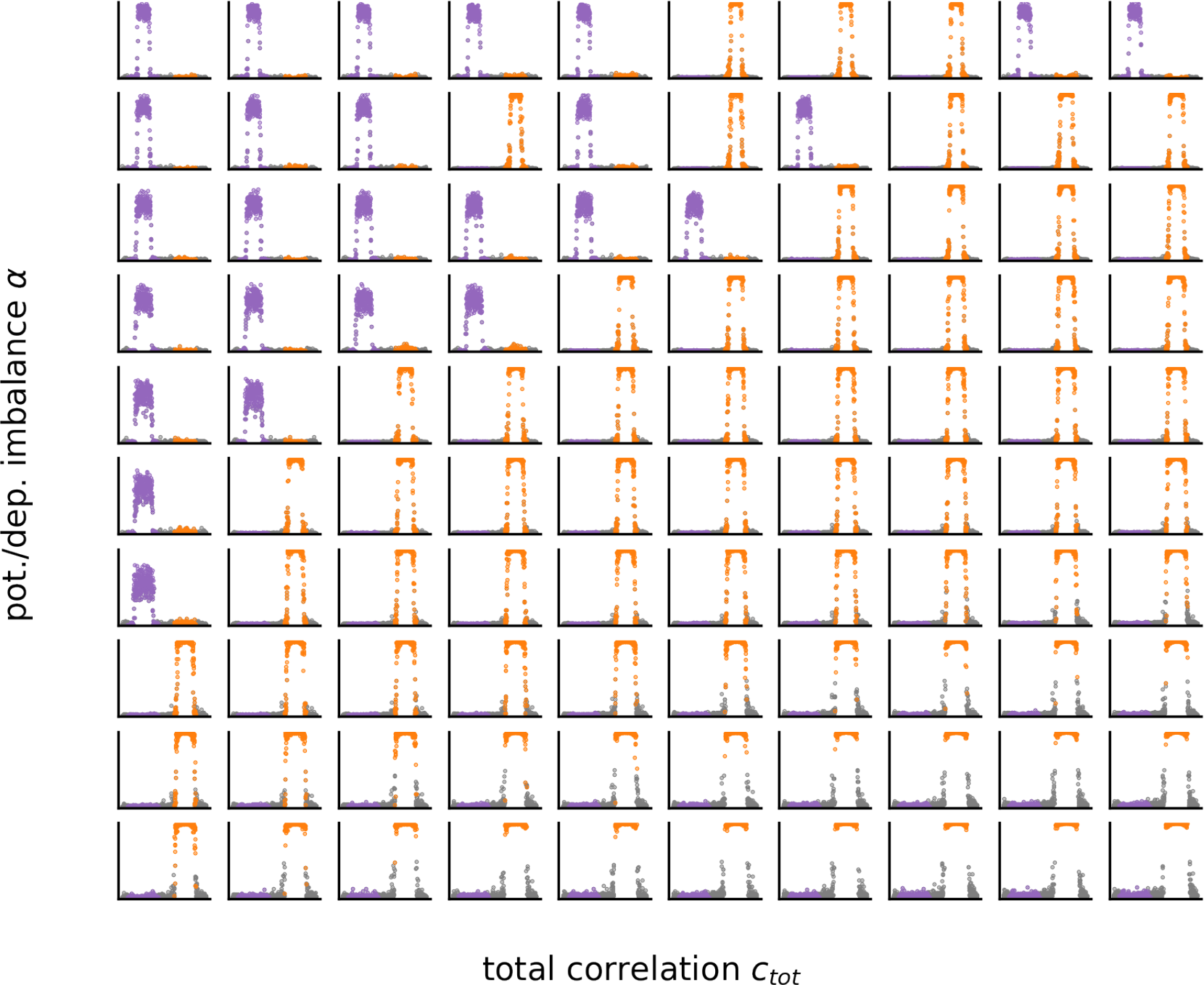
Receptive Fields after convergence for add-STDP (pattern A and then pattern B). Each subpanel corresponds to each of the 10×10 pixels in the heatmaps shown in Figure 4C4.

